# The Drosophila connectome reveals Axo-Axonic Synapses on Descending Neurons

**DOI:** 10.1101/2025.09.04.674108

**Authors:** Cesar Ceballos, Juan Lopez, Ty Roachford, Daniel Sanchez, Sabrina Jara, Kelli Robbins, Casey L Spencer, Rodney Murphey, Rodrigo FO Pena

**Affiliations:** Department of Biological Sciences, Florida Atlantic University, FL 33458, Jupiter, Florida, USA; Harriet Wilkes Honors College, Florida Atlantic University, Jupiter, FL, 33458, USA

**Keywords:** Connectome, Drosophila, Axo-axonic, Giant Fiber escape circuit, Network topology, Electron-microscopy reconstruction, Motor control

## Abstract

Axo-axonic synapses can veto, amplify, or synchronize spikes, yet their circuit-scale logic is unknown. Using the complete electron-microscopy connectome of the adult male Drosophila, we charted every axo-axonic input onto the 1,314 descending neurons that carry brain commands to the ventral nerve cord. By definition, any synapse connected to a descending neuron within the cord is axo-axonic. Thus, we uncovered the ascending-descending and interneurons-descending axo-axonic relationship. Neurons with many partners (high-degree) integrate into the network without clustering into an interconnected ‘rich-club’ of hubs. We identified an octet of ascending neurons (AN08B098) whose axo-axonic input to the Giant Fibers (DNp01) predicted modulation of the escape circuit. Immunostaining confirms their cholinergic identity, while optogenetic activation confirmed that this excitatory cohort increases DNp01 excitability, validating connectome-derived rules. Our work delivers a map of axo-axonic wiring in a complete ventral nerve cord connectome and provides constraints for models of fast motor control.

## Introduction

Neurons communicate through a variety of synaptic motifs. In flies, the canonical arrangement is axo-dendritic, but a second configuration, axo-axonic, can exert unusually powerful control over spike initiation and transmitter release^1,2^. Axo-axonic contacts (AACs) in invertebrates have been known for some time^2–6^. Similar studies have also been conducted in vertebrates^7–9^. In the mammalian cortex, exemplified by chandelier cells, recent studies show how inhibition modulates the pyramidal axons at their initial segments^10,11^. In fish, excitatory spiral fibers and M-cells control a similar circuit^12^. Although these examples suggest broader significance, the quantitative wiring logic of axo-axonic synapses remained poorly characterized, primarily due to the absence of a fully mapped system at the necessary synaptic resolution.

This limitation has been addressed by recent electron microscopy reconstructions of the complete adult male nerve cord and female brain (MANC^13–15^ and FAFB^16^) in *Drosophila melanogaster*, now publicly available through the NeuPrint interface^17^. Early graph analyses of the FAFB data revealed rich-club hubs and other mesoscale motifs^16^, but systematic exploration of the ventral nerve cord (VNC), the insect analogue of the vertebrate spinal cord, is only beginning. At the heart of the VNC, a pair of descending neurons, known as Giant Fibers (GFs), function as command-like neurons that receive visual input in the brain and drive jump and flight motor neurons ^18–20^ Click or tap here to enter text.. These GFs receive a high number of AACs; thus, it is only natural to assume that an AAC-to-behavior modulation is in place. This system is comparable to the descending escape circuits in larger insects, which have been shown to be the target of axo-axonic connections, exemplified by the input to the descending contralateral movement detector (DCMD) neurons in the locust central nervous system^4^.

Within the VNC, a single class of neurons is ideally positioned for studying axo-axonic wiring and their connection to the GFs: Descending neurons (DNs). Because DNs have their somata and dendrites in the brain yet project their axons into the VNC^21–24^, any synapse made onto a DN within the cord is, by definition, axo-axonic. In fact, we have manually curated 1,000 ascending-descending neuron connections, where we observed clear presynaptic mitochondrial expression (93%), a hallmark of axonal terminals^25^ (example shown in Fig. S1, and curation details in Table S1). The MANC connectome contains DN, ascending neurons (ANs), intrinsic interneurons (INs), and motor neurons (MNs), offering a comprehensive substrate for network-wide analysis of this motif. Although classic anatomical work suggested that each GF acts largely in isolation, our work using the MANC connectome reveals an unexpectedly rich set of axo-axonic inputs onto the GF axons, including direct contacts from other DNs, ANs, and INs. In the present study, we show that the GF receives 97 distinct axo-axonic partners, with some offering a crucial role for axo-axonic modulation, i.e., an octet of ascending neurons (AN08B098). Below, we will refer to these neurons of interest as Axc.

Past light-microscopic surveys catalogued individual DN types and noted scattered DN-to-DN contacts^21^, yet the circuit-level role of axo-axonic synapses has remained unexplored. The outstanding issues are their prevalence, partner preference, and functional impact on fast behavior. Specifically, it is unknown how frequently axo-axonic connections occur among the 861,591 possible DN pairs, and whether such a sparse motif can nevertheless mediate rapid, system-wide control required for escape. We tackle these questions by mining the complete electron-microscopy connectome of the adult male ventral nerve cord (MANC). Our analysis (i) enumerates every axo-axonic input onto all 1,314 DNs, (ii) dissects the morphological relationship to DN connectivity, (iii) shows that the DN network is small-world but not rich-club, partitioned into thirteen functional communities, (iv) uses large-scale leaky integrate-and-fire simulations to predict which highly connected AACs modulate the Giant Fiber escape pathway, and (v) confirms those predictions with targeted optogenetic activation *in vivo*.

Our results provide the first system-level blueprint of axo-axonic wiring in the *Drosophila* adult nervous system, suggesting general design principles for descending motor control that may extend beyond insects. They also illustrate the functional power of the sparse axo-axonic network described below and place the GFs at the center of a distributed control system rather than a simple linear reflex arc.

## Results

### Global Distribution of Axo-Axonic Inputs to Descending Neurons

To uncover the prevalence of axo-axonic connections made onto DNs in the VNC, we used the MANC dataset available through the NeuPrint+ interface and API^26–28^. Figure 1*A*-*C* shows morphological reconstructions of some exemplary DNs of interest from Neuroglancer: DNp01 (GFs, black dot), DNd03 (blue dot), and DNg02 (red dot, Fig. 1*A*-*C*). For each neuron, Neuroglancer overlays the 3D-EM reconstruction on the VNC volume. The GFs, whose axons are nearly unbranched (green or blue), receive one of the highest numbers of axo-axonic inputs (n = 97; see NeuPrint+) in the dataset. DNd03 is a highly arborized neuron with a similarly large input count (n=160), while DNg02 has less arborization but similar axo-axonic inputs (n=110). Interestingly, all three share similar synaptic strength (i.e. the equivalent of synapse count from the connectome MANC) despite differences in arborization (DNp01 2352, DNd03 2364, DNg02 1751; Fig. 1*H*). Neuroglancer presents transmission-EM micrographs at 500 nm resolution taken from MANC; presynaptic active zones are marked by magenta dots, postsynaptic densities by cyan dots, and partner axons are pseudo-colored to match the 3D rendering (Fig. 1*D*-*F*).

**Figure 1.**
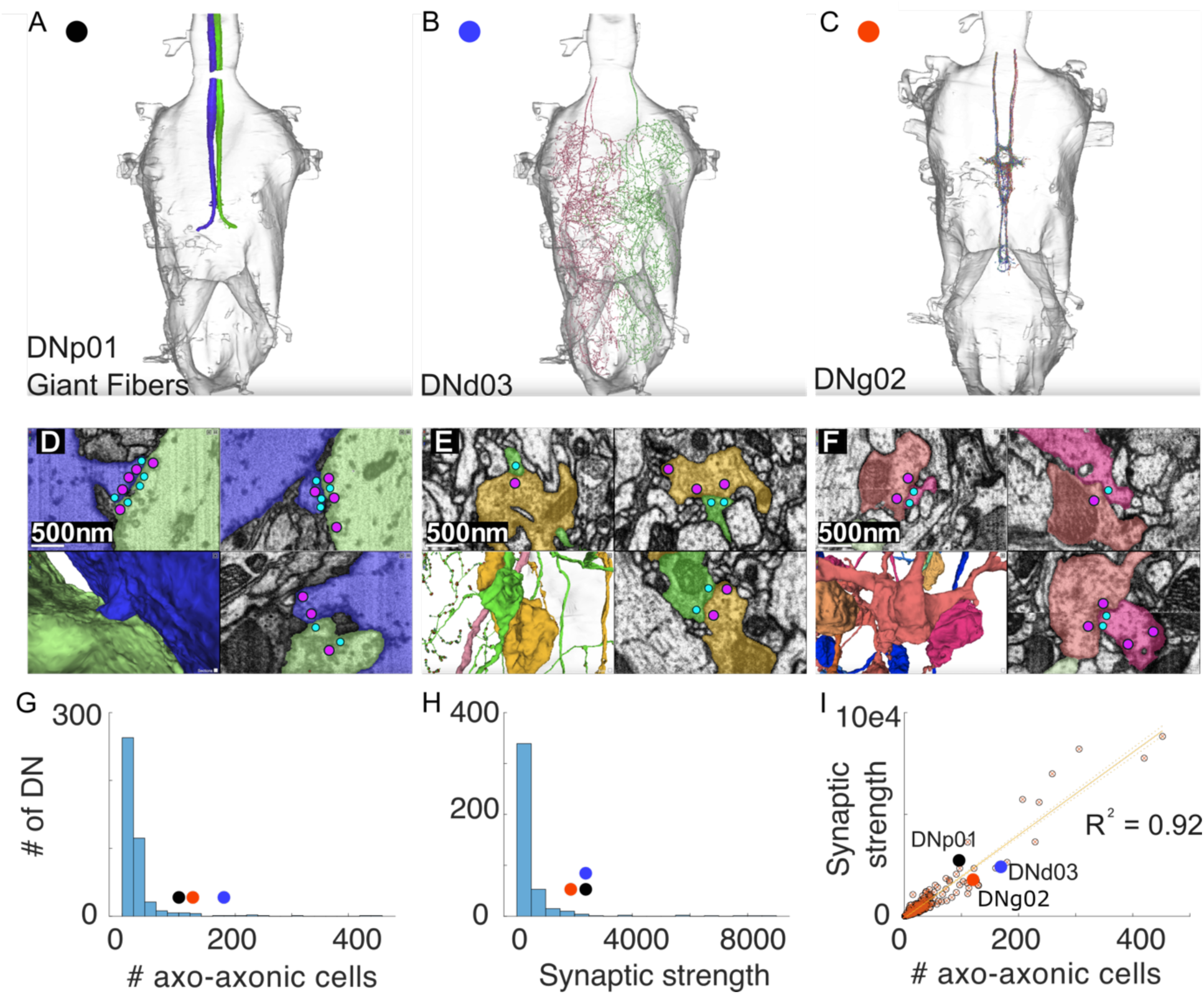
Descending neurons (DNs) and their axo-axonic inputs in the VNC. *(A*)-(*C*) Exemplary Neuroglancer reconstructions of three DNs of interest. (*D*)*-*(*F*) EM images illustrate axo-axonic connections formed by the three reconstructed neurons: (*D*) shows DNp01 to DNp01 connectivity, (*E*) shows DNd03 connectivity with DNp42, and *(F*) shows DNg02 to DNg02 connectivity (same colors as in A-C). Red circles identify the locations of presynaptic T- bars; blue circles show postsynaptic densities. Circles were enlarged for visibility. (*G*) Histogram of the number of axo-axonic neurons innervating one DN. Presynaptic neurons can be DN, AN, or IN. (*H*) Histogram of the cumulative strength of the synaptic connections for each DN. Presynaptic neurons can be DN, AN, or IN. (*I*) Scatter plot and linear regression of the number of axo-axonic neurons vs their synaptic strength. R^2^ = 0.92 and *p <*0.001.

We quantified the distribution of axo-axonic connections (i.e. presynaptic DN, AN and IN) made onto DNs reported by MANC. Figure 1*G* reveals a highly skewed distribution of partner count; the median DN is innervated by a single presynaptic neuron (inter-quartile range 1–3), and 11% of DNs receive input from only one cell with a synaptic strength of six or greater. A small set of outliers, however, attracts hundreds of partners, including DNp01 (GFs) with 97 distinct axo-axonic inputs. Strikingly, the GFs receive these connections despite possessing one of the simplest axonal arbors in the dataset (Fig. 1*A*). All other DNs exhibit more extensively branched and structurally complex axonal arbors, which are likely to account for their higher levels of innervation. Total input strength follows a similar heavy-tailed pattern (Fig. 1*H*). The modal DN receives fewer than 20 synapses, whereas the GF is again among the top of the list in terms of individual contacts. Across the population, partner count and cumulative strength are tightly coupled (Pearson r = 0.96, p < 0.001; Fig. 1*I*), indicating that neurons that recruit many partners also receive proportionally stronger innervation. Together, these findings identify the distribution of axo-axonic connections made with DNs in the VNC. We confirm that the synapses quantified in the graph analysis correspond to active zones and demonstrate how neurons with markedly different morphologies can converge on similar axo-axonic input profiles.

Using these findings, we next examine whether anatomical complexity serves as a strong, though not perfect, predictor of synaptic strength for axo-axonic inputs onto DNs (Fig. 2). Representative reconstructions illustrate the general pattern: neurons with short, unbranched axons receive few contacts, those with intermediate arborization receive proportionately more, and highly branched cells accumulate the greatest axo-axonic strength (Fig. 2*A*-*C*). A kernel density estimation (KDE) plot of all 1,314 DNs reinforces this observation, showing a diagonal ridge in which increasing cable length accompanies increasing total input strength (Fig. 2*F*). Linear, quadratic, and symbolic regressions all yield significant positive slopes, confirming a monotonic relationship (Fig. 2*G*). However, the shallow gradient and the presence of two striking outliers, a long axon with sparse innervation and a short axon with dense innervation, indicate that branch length alone does not determine synaptic investment (Fig. 2*D*-*E*). Thus, we find that a presynaptic neuron’s morphology broadly predicts the degree of modulation received by a partner DN, although additional cell-type-specific factors likely refine the final allocation of axo-axonic synapses.

**Figure 2.**
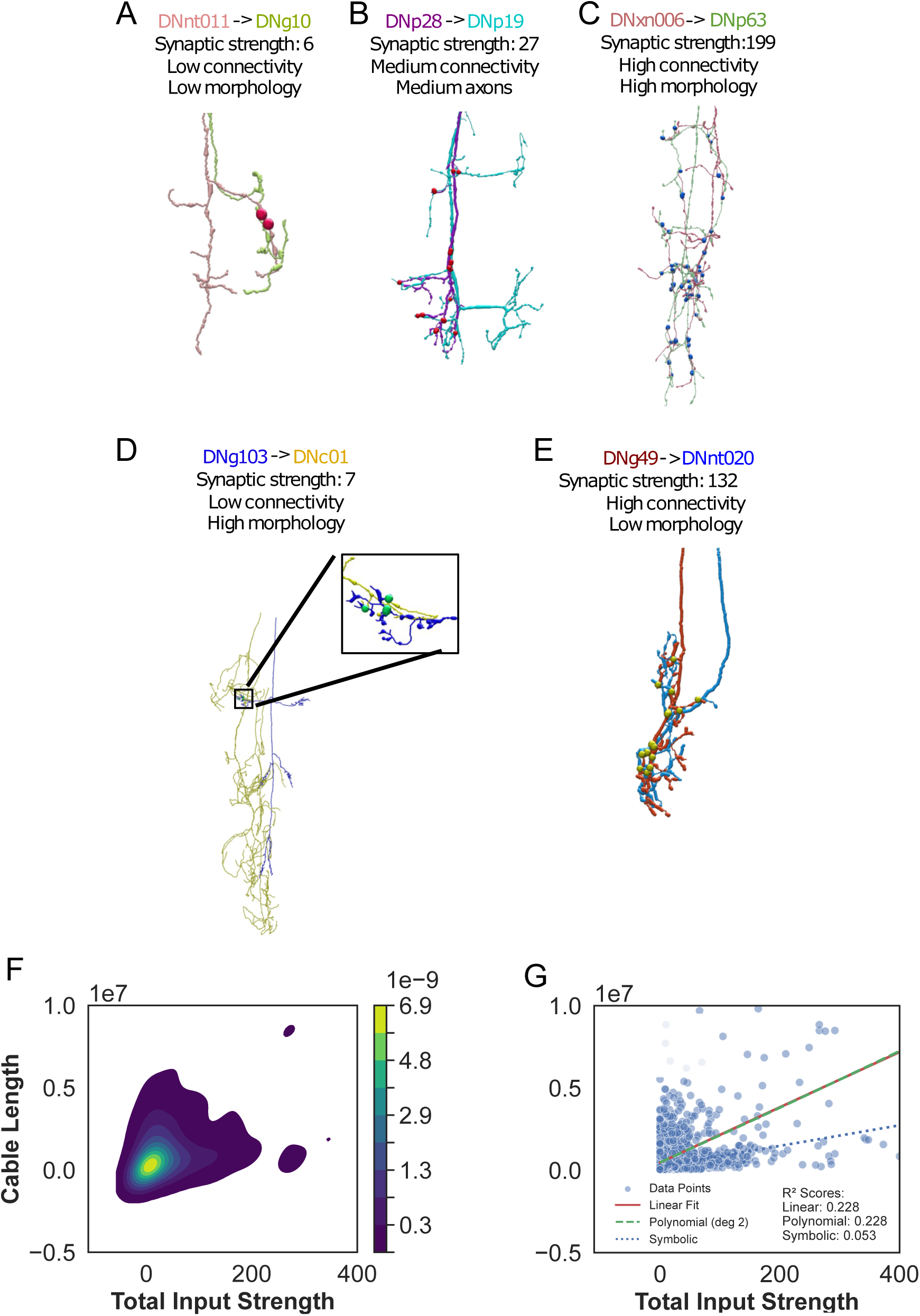
Analysis of morphology and synaptic strength between DN-to-DN axo-axonic connections. (*A*-*E*) Examples of DNs include (legend colors indicate pre- and postsynaptic in the figure): (A) low connectivity with simple morphology, (B) moderate connectivity and morphology, and (C) high connectivity with complex morphology.

Additionally, (D) low connectivity with complex morphology and (E) high connectivity with simple morphology represent exceptions. These neurons generally align with the overall trend—a linear correlation between morphological complexity and synaptic strength—while (D) and (E) are outliers. Dots indicate synapse locations. *F*) Kernel density estimation (KDE) of the scatter data of total input strength vs cable length, *G*) Scatter plot of total input strength vs cable length and the corresponding linear regression, polynomial regression of a quadratic function, and symbolic regression.

Further exploration revealed a unique feature of connectivity. In the VNC, neurons will often form synapses with multiple downstream partners (Fig. 3*A*-*B*). We found certain neurons, such as members of AN08B098, had a disproportionately larger number of connections made onto a single target type of DNs such as the GF than other neuron type. To determine whether these neurons constitute a separable class, we quantified neuron output specialization for every presynaptic neuron using three metrics: the fraction of total synapses made onto the dominant DN, the entropy of the neuron type distribution, and the number of connections of the second largest neuron type connection (off-target) relative to the largest DN connection (on-target).

**Figure 3.**
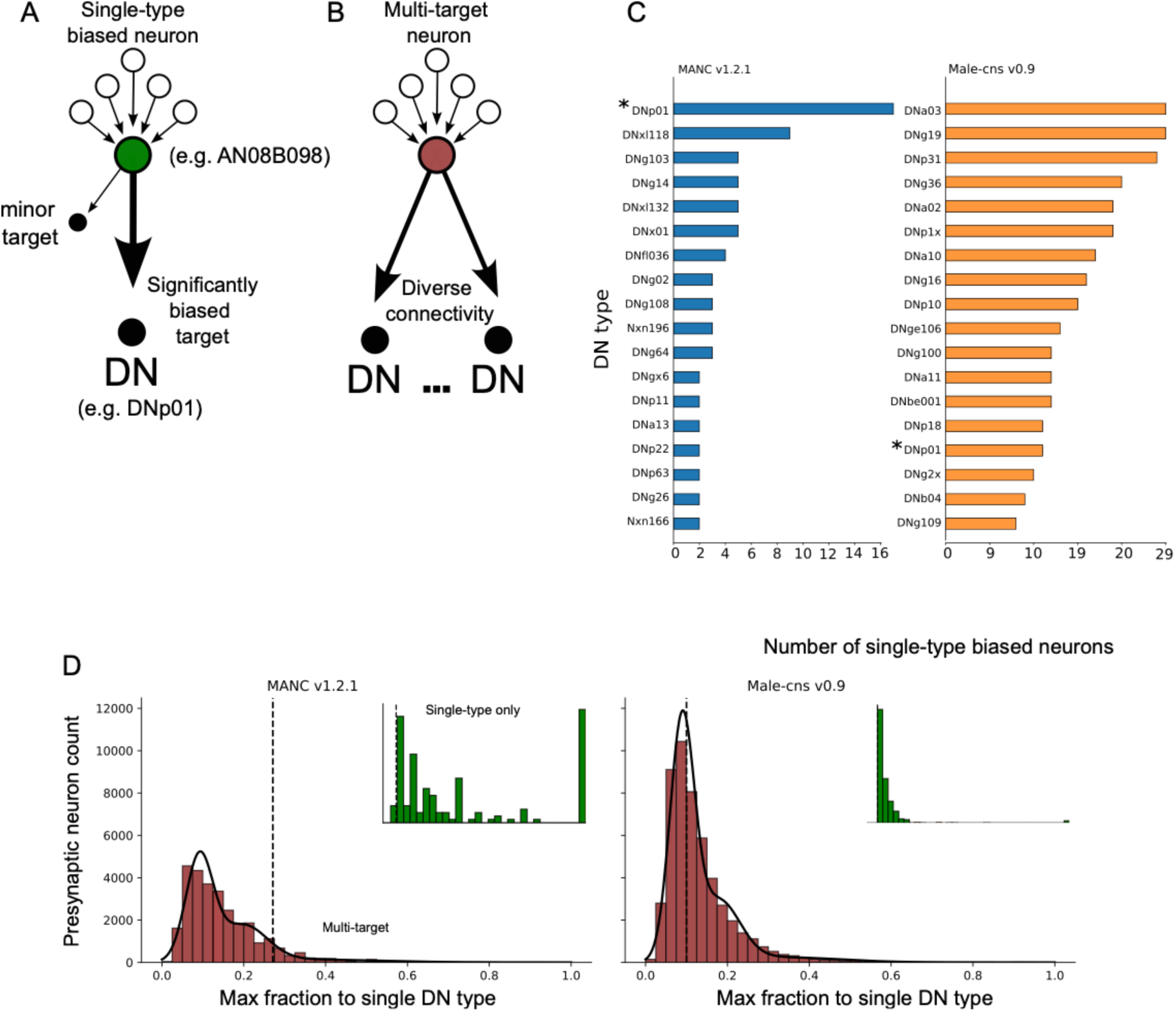
Neurons that provide exclusive axo-axonic connections to DNs. (*A*) Sketch of single type biased vs (*B*) multi-target neuron. Single-type biased neurons project a significant proportion of their outgoing connections to a single DN, while unbiased neurons form several strong connections among partners. (*C*) Number of exclusive neurons per DN for both MANC v1.2.1 (blue histogram) and Male-cns v0.9 (orange histogram). The Giant Fibers (DNp01; marked by an asterisk) are among a small population of DNs that receive most single-type biased connections, i.e., as seen in (D), there are approximately 16 neurons that innervate DNp01 in MANC and 11 in Male-cns. (*D*) For each dataset, the fraction of outputs made from neurons onto their most strongly targeted DN type was computed per neuron, and a dataset-specific cutoff was identified from the distribution as the point separating the population of unbiased neurons and single-type biased neurons (MANC 0.27, male-cns 0.1). Single-type biased neurons across both datasets had similar properties, including high target preference, reduced target entropy, and reduced off-target connection strength (Kolmogorov-Smirnov test, p < 0.001). We further refined this by removing neurons whose off- target connections had 15 or greater synapses, suggesting non-influence over their synaptic partners^29^.

For each presynaptic neuron, we first quantified the percentage of strength of its outgoing synapses that were made onto its most strongly targeted DN. Neurons were required to exceed a threshold value and the percentage of strength of the dominant DN was permitted to exceed the threshold, ensuring a uniquely dominant target. Initially, we assumed threshold values that separate two populations: multi-target vs single-type biases. The final threshold was determined by optimizing the minimization of an error parameter. A statistical comparison of the distributions using the Kolmogorov-Smirnov test (with a significant p-value < 0.001) revealed two distinct distributions of presynaptic neurons, which we defined as single-type biased and multi-target, with threshold values of 0.27 in MANC v1.2.1 and 0.1 in male-cns v0.9. In addition, single-biased cannot send more than 15 connections to other off-target DN to exclude external influence^29^.

In both connectomes, we identified several single-type biased axo-axonic connections (Fig. 3*C*) to DNs in both MANC v1.2.1 and male-cns v0.9. DNp01 receives the highest number of single-target AACs in MANC (n = 16) and among the highest for male-cns (n = 11). Among these neurons, five belong to the Axc type, and the remaining are labeled as “unknown”.

Compared with other highly connected DNs, the GFs attain an equivalent synaptic strength with a simple morphology, further demonstrating how morphological complexity alone does not predict axo-axonic input. These results identify the GFs as exceptional hubs for axo-axonic modulation due to their significant connectivity despite their relatively simple morphology compared to other highly connected DNs. However, these findings raise several new questions: How are AAC populations connected, and how do they act on GFs? Is the GF an outlier, or do similar axo-axonic motifs govern other rapid motor pathways in the fly? How do these contacts alter GF excitability? We address these questions by investigating the strongest single-target AAC population presynaptic to the GFs, the Axc type neurons.

### Sparse yet Globally Connected Axo-Axonic Network

To address how AAC populations are connected, and their influence on other DNs, we extracted every DN-to-DN contact from the updated MANC dataset. The resulting adjacency matrix is strikingly sparse: only about 1% of the possible ordered pairs carry even a single synapse (Fig. 4*A*). Moreover, partner choice is not random: homotypic pairs (same DN type) and heterotypic pairs (different types) show distinct transmitter profiles, as shown for acetylcholine, GABA, and glutamate, implying that neurotransmitter identity contributes to the selectivity of these long-range axo-axonic contacts (Fig. 4*D*). In homotypic DN connections, ∼90% of the connections are cholinergic (excitatory), while heterotypic DN connections are ∼60% cholinergic [the remaining are GABA/glutamate (inhibitory)]. This suggests that AACs between DNs, while rare, are predominantly excitatory. connections.

**Figure 4.**
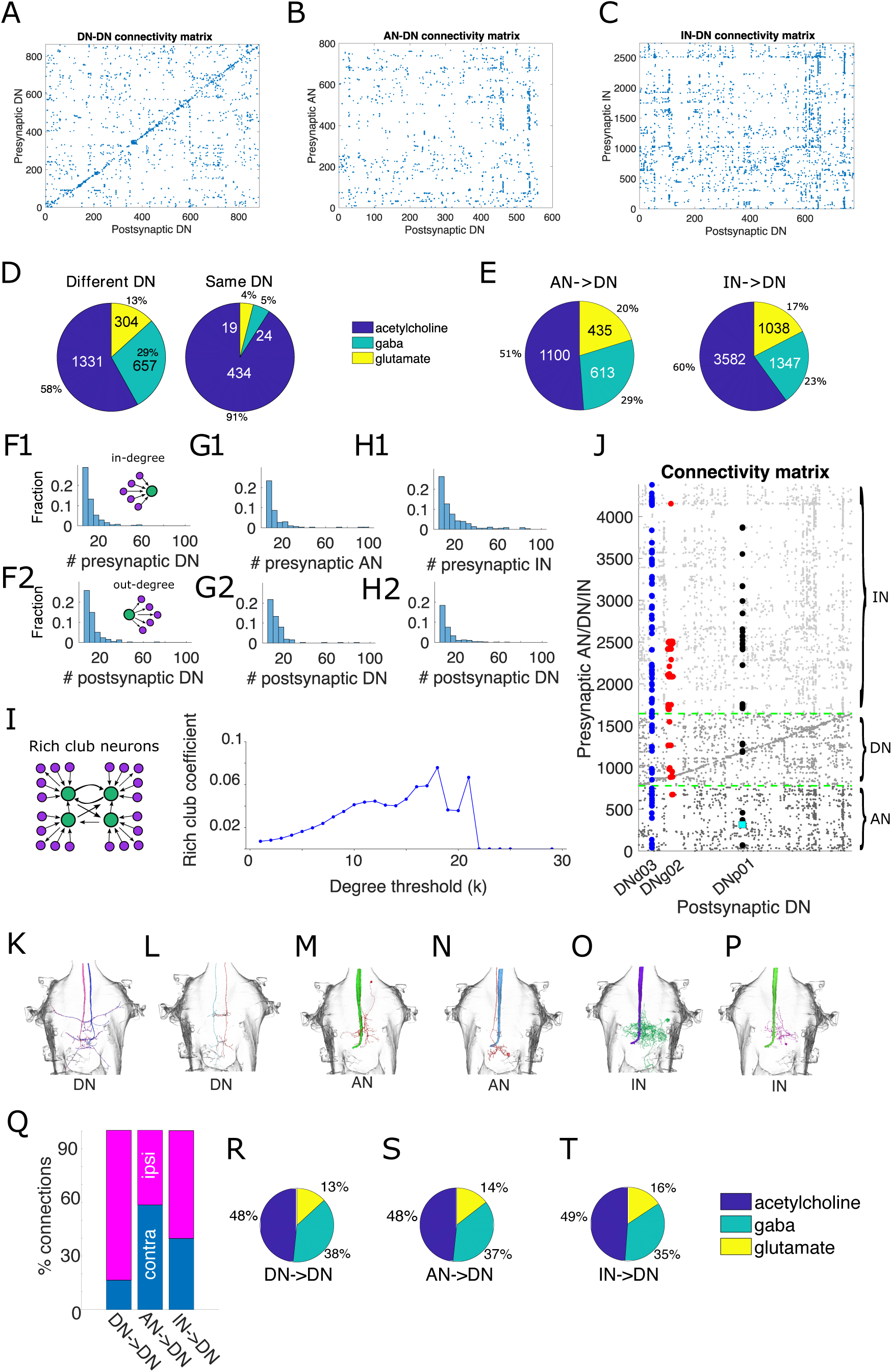
Axo-axonic connectivity analysis. (*A*) Connectivity matrix of pairs of DN-to-DN pairs. Only 1% of all possible connections were identified, suggesting that DN-to-DN axo-axonic connections are extremely rare. (*B*) Connectivity matrix of pairs of AN-to-DN pairs. Only 1.2% of all possible connections were found, which suggests that AN-to-DN axo-axonic connections are also exceptionally rare. (*C*) Connectivity matrix of pairs of IN-to-DN pairs. Only 0.7% of all possible connections were observed, indicating that IN-to-DN axo-axonic connections are uncommon. (*D*) Pie chart of predicted neurotransmitter of DN of the different type (left, e.g., DNp01 -> DNp02) and same type (right, e.g., DNp01 ->DNp01) that make pairs of axo-axonic connections. (*E*) Pie chart of predicted neurotransmitter of AN-to-DN and IN-to-DN axo-axonic connections. (*F*) Histograms of the number of (F1) presynaptic DN (in-degree) and (F2) postsynaptic DN (out-degree) for DN-to-DN pairs. (*G*) Histograms of the number of (G1) presynaptic AN and (G2) postsynaptic DN for AN-to-DN pairs. (*H*) Histograms of the number of (H1) presynaptic IN and (H2) postsynaptic DN for IN-to-DN pairs. (*I*) Rich-club analysis of the DN-to-DN axo-axonic network. (*J*) Connectivity matrix of pairs of AN/DN/IN-to-DN connections. Filled circles follow the same color scheme as in Fig. 1A-*C*. Cyan square identifies Axc-type neurons. Dashed green lines and gray tones for dots separate ANs, DNs, and INs. (*K*-*P*) Examples of each category from DN-to-DN, AN-to-DN, and IN-to-DN. (From left to right: (*K*) DNx01-DNx01, (*L*) DNb04-DNb04, (*M*) AN08B099-DNp01, (*N*) AN10B019-DNp01, (*O*) IN11A001-DNp01 and (*P*) IN06B059-DNp01). (*Q*) Percentage of contralateral and ipsilateral connections between DN-to-DN, AN-to-DN, and IN-to-DN. (*R*-*T*) Pie chart of predicted neurotransmitter of DN to contralateral DN, AN to contralateral DN, and IN to contralateral DN of axo-axonic connections.

We then asked whether the same wiring principle applies to non-descending inputs, namely ascending neurons (ANs) and intrinsic interneurons (INs). Repeating the analysis with ANs and INs as presynaptic partners to DNs again produced very sparse matrices: about 1.2% of all possible AN-to-DN pairs and 0.7% of the IN-to-DN pairs contain at least one synapse (Fig. 4*B*-*C*). Figure 4*E* shows the neurotransmitter profile for AN-to-DN and IN-to-DN pairs. Both pair profiles were similar, with nearly half of the neurons classified as cholinergic and the other half GABA/glutamate. This suggests that AACs between AN-to-DN and IN-to-DN follow the same rules for neurotransmitter identity as DN-to-DN AACs.

Lastly, we show the distribution of the number of pre-synaptic DN, AN, or IN that project axo-axonic synapses to a post-synaptic DN (Fig. 4*F*-*H* top row) as well as the number of postsynaptic DN that are innervated axo-axonically by the same DN, AN, or IN (Fig. 4*F*-*H* bottom row). The distributions are similar across neuron types, showing a decay pattern resembling an exponential (Poisson) distribution, where a few neurons are highly connected, and a few neurons are sparsely connected. When we merge all three presynaptic classes into a single connectivity matrix (Fig. 4*J*,) the same pattern from Figure 4*A*-*C* emerges: extreme sparseness (see dots where our exemplary neurons are located; color scheme as in Fig. 1*A*.). Note that in Fig. 4*A* there is a clear diagonal line which indicates DNs synapsing onto their contralateral counterparts. We also highlight the degree of convergence across presynaptic classes and identify connectivity hubs based on their total input. Taken together, regardless of presynaptic identity, the findings demonstrate that the network’s organization combines stringent partner selectivity with complete coverage of the descending system.

Global wiring principles were identified to explore more refined AAC spatial connectivity preferences and determine whether they are repeated across DNs, ANs, and INs. Figure 4 shows examples of the contralateral axo-axonic connections between DN-to-DN (Fig. 4*K-L*), AN-to-DN (Fig. 4*M-N*), and IN-to-DN (Fig. 4*O-P*). Figure 4*Q* shows the percentage of contralateral and ipsilateral connections. AACs between contralateral DNs are rare. However, ANs and INs projecting to contralateral DNs are frequent. Interestingly, the neurotransmitter profiles from our earlier findings are preserved regardless of the type of presynaptic neuron, where approximately half of the neurons are cholinergic (excitatory), and the other half is GABAergic/glutamatergic (inhibitory) (Fig. 4*R*-*T*).

Collectively, these findings suggest a coherent narrative. Whether the presynaptic axon descends, ascends, or remains intrinsic to the cord, axo-axonic wiring obeys the same two rules: partner selectivity is extremely high, and midline crossing is common only for neurons that share strong reciprocal ties. The result is a VNC in which local subtypes can interact tightly with their mirror images, yet the great majority of neurons remain insulated from direct axo-axonic influence.

### Graph-Theoretic Architecture of the Axo-Axonic Network

To address if GF is an outlier or do similar axo-axonic motifs govern other rapid motor pathways in the fly, we study connectivity matrices that are derived employed as a directed graph to examine the system-wide arrangement of sparse axo-axonic interactions among DNs (Fig. 4*A*). The graph contains 884 nodes and 2,757 edges, giving a density of 0.007. Despite this low density, the average shortest-path length is only 4.03, indicating that any two DNs can be reached through a small number of intermediate axonal contacts. In fact, we can compare with the chance level: the formula ⟨*L*⟩ = ln(*n*) /ln (*k*^−^) is a well-known and widely used approximation for the average shortest-path length in large random networks, such as the Erdős-Rényi model. Here, *n* is the number of vertices in the graph, *k*^−^ is the mean degree, which can be obtained by the ratio *m*/*n* with *m* being the edge count. In our problem, *n* = 884 DNs and *m* = 2,757 axo-axonic contacts leave us with ⟨*L*⟩ ≈ 5.9. Therefore, the average shortest path length is indeed shorter than you would expect by chance. The clustering coefficient is likewise low at 0.013, indicating limited local redundancy. Community detection identifies 13 separate modules, consistent with the network’s functional subdivision. Node degree ranges from 1 to 29 with a mean of 6.24, reflecting a modest hierarchical spread.

The rich-club coefficient Φ(k) measures the density of connections among nodes whose total degree (in-degree + out-degree) equals or exceeds a threshold k. Φ(k) = 1 indicates that the high-degree subgraph is fully connected, whereas Φ(k) = 0 means that those nodes share no edges beyond chance level. The rich-club coefficient in our data peaks at 0.076 for nodes with a degree of at least 18, so high-degree hubs do not preferentially connect to each other; integration is therefore distributed rather than centralized (Fig. 4*I*). The modest peak value and the absence of a sustained plateau indicate that high-degree DNs do not preferentially interconnect to form a rich club; instead, network integration is distributed rather than hub-dominated.

A slightly positive degree assortativity of 0.029, a measure that suggests a tendency of nodes to connect with other nodes in a similar way, suggests a weak tendency for neurons of similar degree to link. Finally, the maximum betweenness centrality of 0.049, a measure indicating how often a node lies on the shortest path between all pairs of nodes in a network, highlights a small set of bridge neurons that channel information between modules. Together, these metrics portray a network that combines efficient global communication with modular specialization, an architecture well-suited for coordinating diverse motor outputs.

### Axo-axonic modulators of the Giant Fiber escape circuit

The MANC connectome introduces several novel neuron types to the Giant Fiber escape circuit. To understand and underscore the significance of AAC modulation of the circuit, it is important to know which individual cells can exert a strong, targeted influence on a system such as the GF system^18,19^. As shown in Figure 3, the GFs are among a small population of DNs that receive single-target connections. A focused search within MANC revealed the Axc type neurons, a cluster of eight neurons whose axo-axonic output is both nearly exclusive and among the strongest to the GFs (Figs. 3-4). These Axc-type neurons are predicted to be cholinergic by the MANC connectome. The BANC connectome demonstrates that these neurons do not connect to the GF in the brain (see Fig. S3).

To test whether the connectome-predicted Axc type neurons truly form chemical AACs with the GFs, we used split-Gal4 drivers to direct membrane-bound GFP to Axc axons, back-filled the GFs with tetramethyl-rhodamine, and labelled presynaptic active zones with anti-Bruchpilot (nc82). High-resolution confocal stacks revealed discrete GFP-positive boutons that colocalize with Brp puncta on the descending GF tract in the first thoracic neuromere (Fig. 5*A*,*C*), confirming axo-axonic contacts between Axc type neurons and GF. The split-Gal4 line being used labels AN08B098. However, there are a few extra neurons labeled that can come from AN08B099 and make weak connections to the GF. We have also provided (i) single-section confocal images with optical enhancement showing the synaptic connectivity and (ii) similar EM reconstructions from Neuprint to demonstrate that synapse locations along the axon correspond to synapse locations identified in the MANC connectome (Fig.5*G-O*). Immunostaining for Choline Acetyltransferase (ChAT) revealed that Axc somata, as labelled through genetic expression of mCD8::GFP, are ChAT-positive (Fig. 5*D*-*G*), establishing this cohort as likely cholinergic and therefore excitatory.

**Figure 5.**
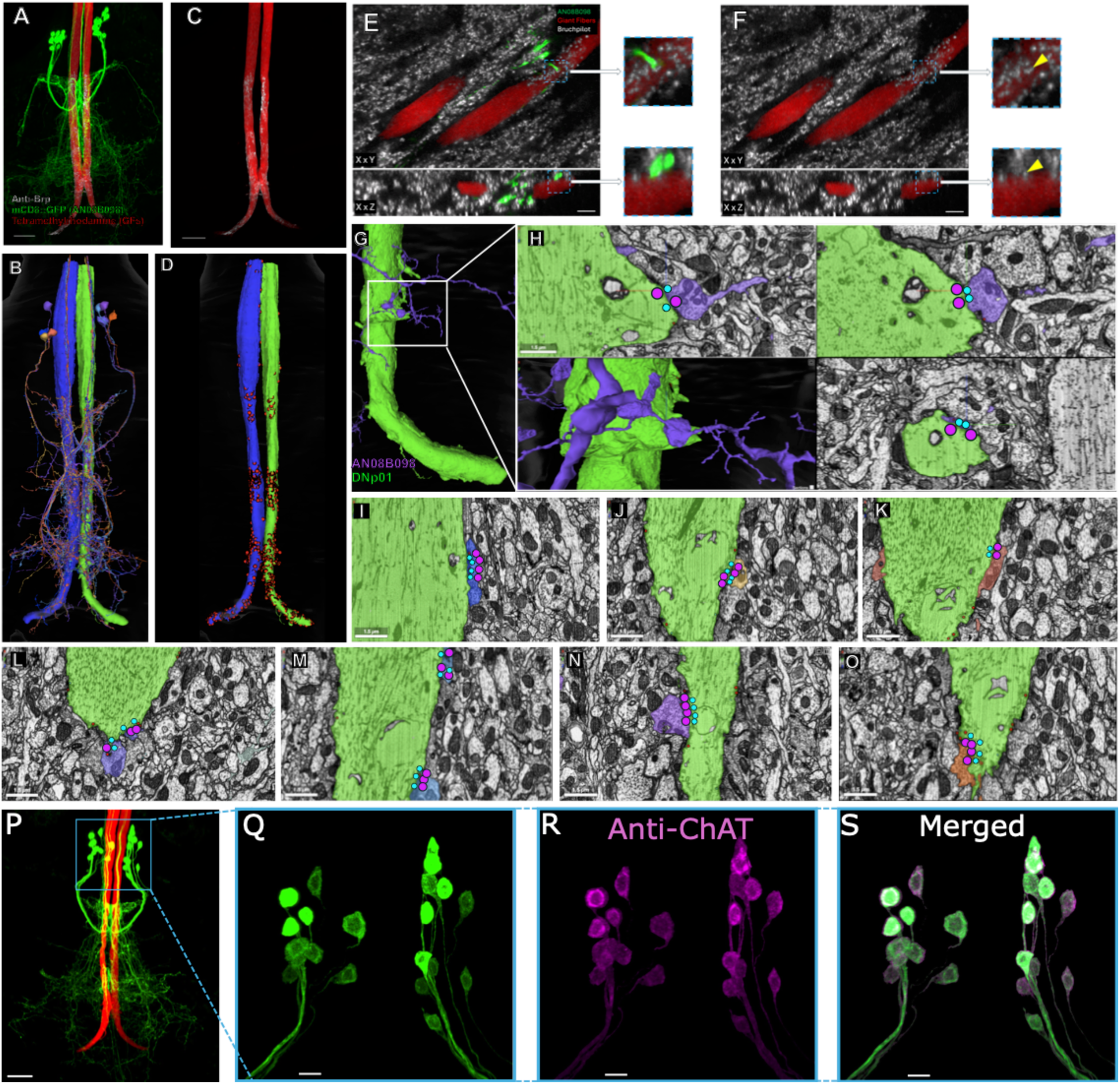
Axcs make cholinergic axo-axonic synapses on the GFs as predicted by the MANC connectome. (*A*) AN08B098-type neurons (green) form axo-axonic connections at distinct locations along Giant Fiber (red, GF) axons. The scale bar shown is 20 µm. (*B*) The AN08B098-GF connections reconstructed in Neuroglancer mirror the morphology seen with fluorescence in *A*. (*C*) We validated AN08B098-to-GF connectivity using anti-bruchpilot to stain for T-bars in active zones (white) along AN08B098-type neurons and colocalized the resulting fluorescence with the GFs (red), which were filled with tetramethylrhodamine. The scale bar shown is 20 µm. (*D*) Neuroglancer reveals presynaptic sites at similar locations seen in colocalized fluorescent image *C*. (*E-F*) XY, and XZ plane view of the preparation shown in *A* and *B*. The zoomed-in 6 µm inlays show AN08B098 forming a single synapse with the GF in E. The yellow arrows detail the precise location AN08B098 forms the Brp-positive chemical synapse (white) to the GFs in F. Scale bars shown are 5 µm. (*G*-*O*) EM images showing monosynaptic connections between single GF (green) and AN08B098 neurons identified by the following MANC id: (G-H) 21041, (I) 21589, (J) 23949, (K) 152261, 16900, (M) 20444, (N) 22275, and (O) 24038. Pre-and postsynaptic sites are detected in EM slices using a 3D convolutional neural network to identify T-bars (cyan dots) and postsynaptic densities^30^ (PSDs, magenta dots). (*P*-*S*) We expressed anti-choline acetyltransferase (anti-ChAT) and anti-GFP in AN08B098 neurons. Anti-ChAT colocalizes to AN08B098 cells, with particularly bright staining in the cell bodies. This finding suggests acetylcholine synthesis is present within these cells, validating connectome transmitter predictions for AN08B098.

To address how these contacts alter GF excitability, and following this imaging confirmation, we investigated whether these AACs could measurably influence GF activity in a simulation of the complete VNC connectome. To test this, we stimulated all VNC neurons with a Poisson distribution of excitatory synaptic currents at 30 Hz and 5 nS. Additionally, one GF was stimulated more strongly at 10 Hz and 40 nS to establish a baseline level of excitability (Fig. 6*A*, *where* spikes are indicated by large black dots at the bottom of the raster plot and arrows underneath). When the eight cholinergic Axc neurons were activated, the GF crossed threshold more often, indicating that the axo-axonic inputs are net excitatory (Fig. 6*B*). Additional simulations with different scenarios where stimulation and deletion of non-Axc (nAxc) populations demonstrate that the Axc cells are paramount for increased excitability (Figs. 6*B*-*E*). GF subthreshold voltage histograms quantify the increased excitability (Fig. 6*F*), supported by statistical significance (Fig. 6*H*). These simulations highlight a previously unrecognized cohort of axo-axonic synaptic modulation, providing a rapid mechanism for amplifying the escape-circuit output.

**Figure 6.**
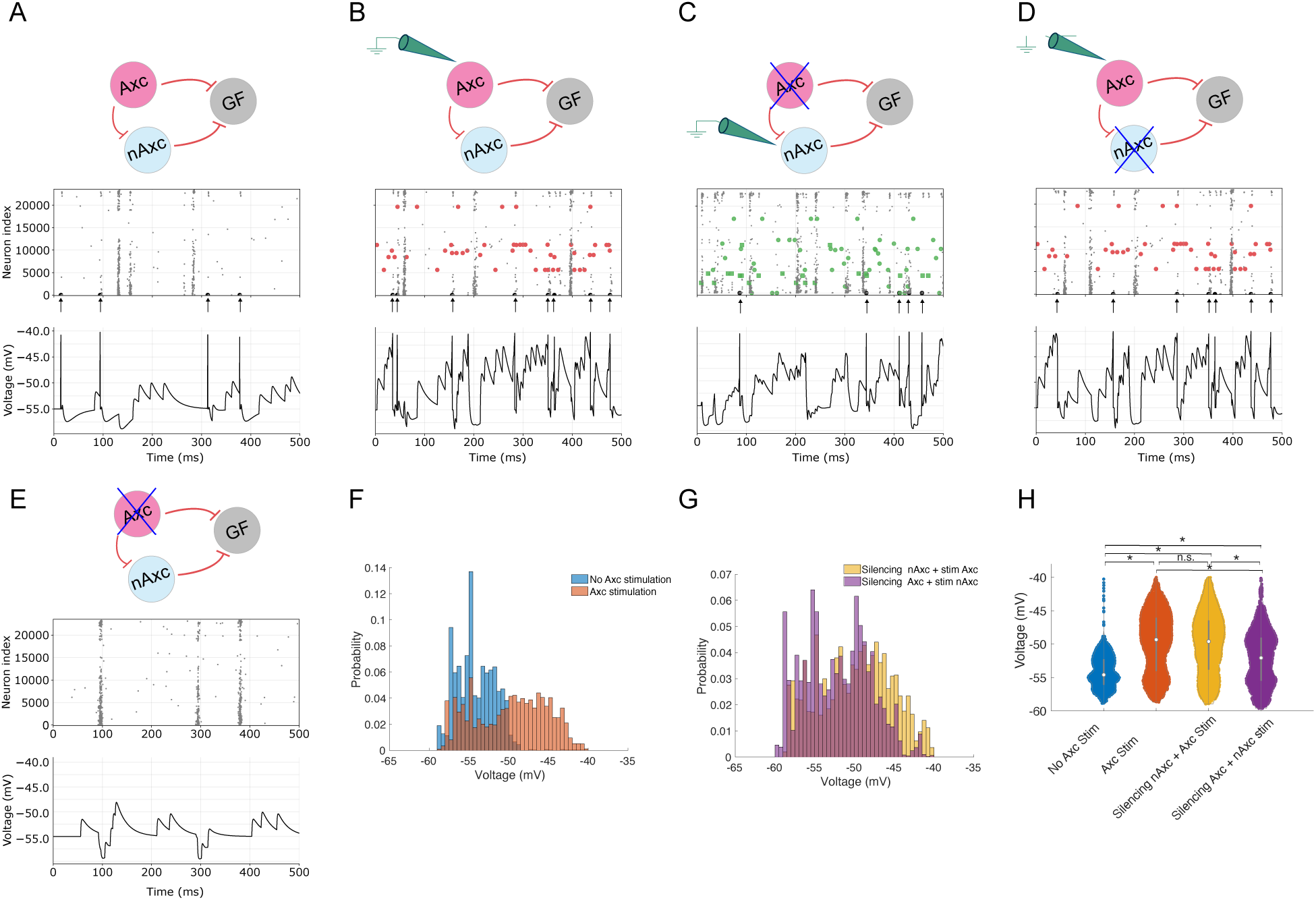
Simulations using the whole VNC network. (*A*) Stimulation of the GF and corresponding raster plot of all the neurons in the VNC and voltage trace of GF response (spikes are shown as large black dots at the bottom of the raster plot, also marked with an arrow). (*B*) Schematic of the GF stimulation by Axc type neurons and corresponding raster plot of all the neurons in the VNC and voltage trace of GF response (GF spikes are represented by large black dots at the bottom of the raster plot, while Axc spikes are represented by large red dots). (*C*) Similar to previous plots, but now the stimulation is driving non-Axc (nAxc), and the Axc is removed from the simulation. (*D*) Similar to previous plots, but now the stimulation is driving Axc cells, and nAxc are deleted. (*E*) Similar to *A*, but now Axc cells are deleted. (*F*) Histogram comparison of subthreshold voltages during GF-only and GF+Axc stimulation. (*G*) Similar to *F*, but now the histogram shows the difference between silencing nAxc and stimulation in Axc vs. silencing Axc and stimulation in nAxc. (*H*) Violin plots showing statistical analysis of some of the comparisons. The overall excitability of the GF increases when the Axc neurons are activated, leading to higher firing probability for the GF.

As the control experiment, we also tested the possibility of silencing nAxc cells, which are defined here as presynaptic cells to the GF that contribute to polysynaptic pathways, and Axc cells selectively (Figs. 6*C*-*D*). The results from these figures demonstrate that when nAxc connections are silenced, the increased excitability persists, and when the Axc monosynaptic connections are silenced, but nAxc is retained, the excitability is decreased. Fig. 6*G*-*H* shows additional quantification of this Axc to GF excitability, with the only non-significant comparison being Axc stim vs. silencing nAxc + Axc stim, demonstrating that the possibility that Axc acts polysynaptically could be neglected.

### Testing connectome predictions *in vivo* via optogenetic stimulation of axo-axonic neurons

To assess the predictions of these connectome simulations, we tested the ideas *in vivo* using classic electrophysiology and optogenetics. We used the split-Gal4 line that labels the Axc neurons (BDSC #89945) to drive a UAS construct containing a mVenus-tagged red-shifted channelrhodopsin, CsChrimson (BDSC #55136). Expression of CsChrimson was verified with immunohistochemistry (Fig. S2A). We electrically stimulated the GFs at 24 Hz in flies expressing CsChrimson in Axc type neurons and recorded from the jump muscle (TTM) and flight muscle (DLM) as previously described^31^, (Fig. 7*A-B*). We then set the electrical stimulation of the GF just below threshold. This near-threshold stimulation resulted in occasional (∼20%) responses in both muscles (n = 5 specimens, Fig. 7*B* and 7*E*). However, when the same preparation was illuminated (590 nm, 0.37mW) during the stimulus train, the sub-threshold electrical stimulus elicited responses from both muscles nearly 100% of the time (n = 5 specimens, Fig. 7*D* and 7F). A one-way ANOVA found significant differences between responses before, during, and after light application (*p <* 0.001), but there was no significant difference for the control group (*p =* 0.53) (Fig. *7G*). Post-hoc multiple comparisons revealed significant differences between responses at baseline (before light application) and during and after light application. We also found significant differences using a paired t-test for the same responses during and after light application (*p<*0.001), but not before (*p>*0.26).

**Figure 7.**
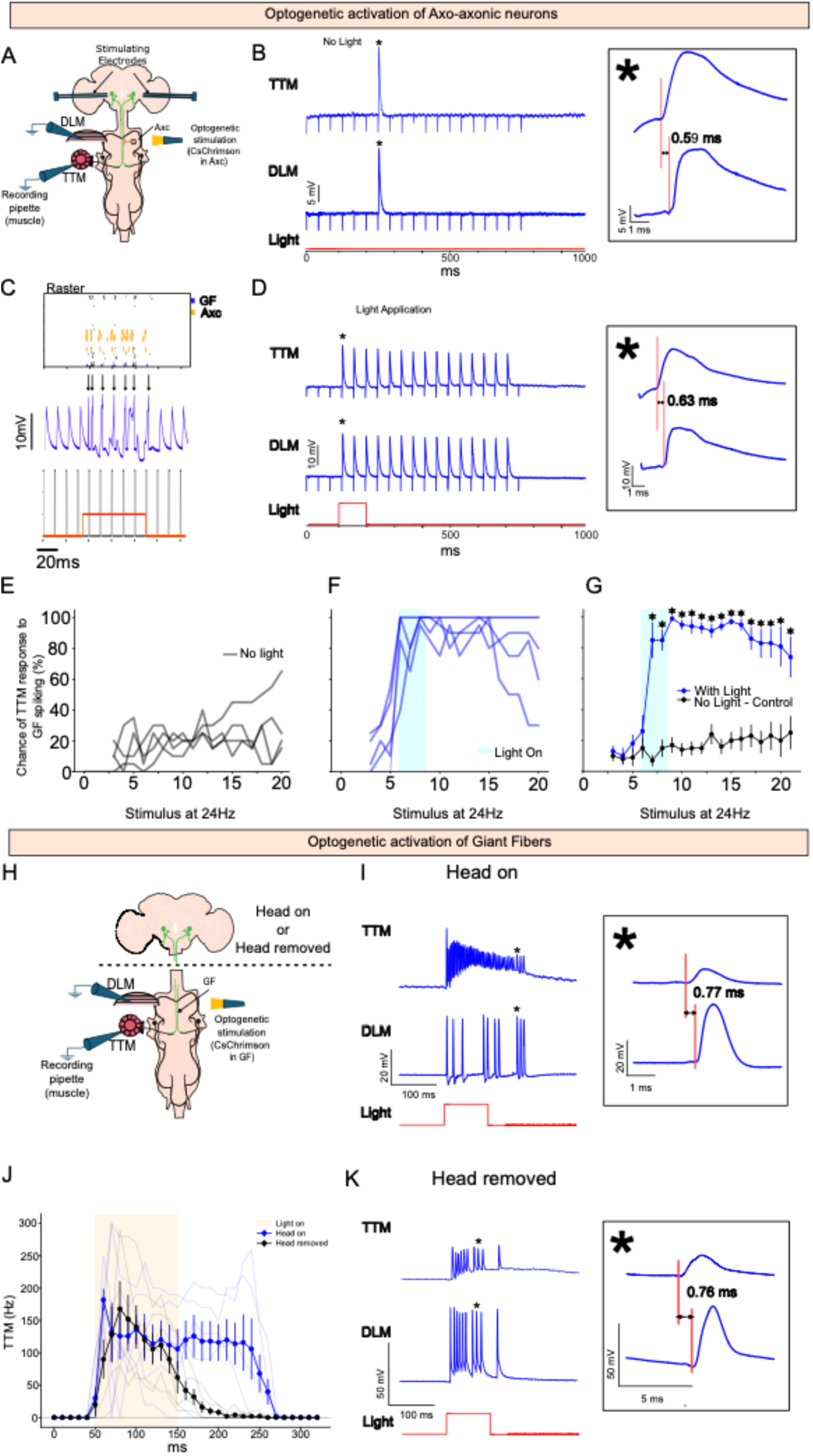
Optogenetic activation of the Giant Fiber system. (*A*) Sketch of the electrophysiological recordings with optogenetic activation of axo-axonic cells (Axc). (*B*) Example of raw traces for control (no light application). A zoomed-in panel is presented, showing a constant difference in latencies of ∼0.6 ms between TTM and DLM indicative of GF spiking. (*C*) Simulation of a computational model of the VNC connectome emulating the experimental results under subthreshold conditions and light stimulation. Note that GF only spikes when the light is applied and is time-locked with the electrical pulses. Action potentials are indicated with arrows. (*D*) In vivo example of raw traces when light was applied. A zoomed-in panel is presented, showing difference in latencies of ∼0.6 ms between TTM and DLM responses. (*E*) Response probability per fly (n = 5) under no-light conditions. (*F*) response probability per fly (n = 5) under light conditions (light was applied during the blue shaded region). (G) Average and SEM of response probability (paired t-test, * = *p*<0.001). **Spike initiation in the VNC**. (*H*) Sketch of the electrophysiological recordings with optogenetic activation of the GF before and after decapitation. (*I*) Recordings from both TTM and DLM before decapitation. The constant latency between TTM and DLM indicates the presence of GF action potentials. A zoomed-in panel shows the constant latency between TTM and DLM ∼0.77 ms. (*J)* Response of the TTM to optical stimulation in a specimen driving CsChrimson in the GF as above. The blue curve represents the response of the TTM in animals with the head on, exhibiting a long tail of post-light response. The black curve is after the head was removed to remove the spike-initiating zone in the brain. (*K*) Recordings from both TTM and DLM after decapitation. The constant latency between TTM and DLM indicates the presence of GF action potentials. A zoomed-in panel shows the constant latency between TTM and DLM ∼0.76 ms.

To verify that light stimulation does not elicit effects in the absence of CsChrimson expression or function, we performed two control experiments. In the first experiment, flies expressed GFP instead of CsChrimson (Fig. S2A); in the second, CsChrimson-expressing flies were raised without dietary cis-trans-retinal, thereby preventing optogenetic activation (Fig. S2*B*). A one-way ANOVA revealed no significant differences in stimulus-evoked responses across conditions (*p* = 1). Post-hoc comparisons revealed no significant differences between the baseline, light, and post-light periods. Paired t-tests also found no significant differences between light and no-light conditions during and after stimulation. These results confirm that the observed effects are specifically attributable to CsChrimson activation.

How the Axc alters excitability depends in part on the site of spike initiation for the GF. In the present experiments, the stimulating electrodes are in the brain where there are no monosynaptic connections from Axc to GF^26^. The only monosynaptic Axc-to-GF connections are in the VNC (Fig. 5, S3). How can the optical excitation of the Axcs in the VNC, where they make synaptic contact, have a modulatory effect on the GF spike initiation zone (SIZ) in the brain, where they do not make contact? We have now demonstrated that the GF also possesses a spike-initiating region(s) in the VNC, where the Axc could influence spike initiation. We demonstrated this alternative SIZ by driving CsChrimson in the GF, removing the head to remove the SIZ in the brain, and testing headless animals with optical stimulation to reveal GF activity generated in the VNC (Fig. 7J-*K*). GF activity driven by the light stimulation is revealed by the response of both TTM and DLM with a constant latency between them indicative of GF action potentials (Fig. 7*K*). Therefore, the Axcs can directly influence this SIZ in the VNC.

Unexpectedly, we also note a prolonged response of the GF after the light was turned off (Fig. 7D, 7F and 7G). Before light application, the TTMn response probability was low (∼20%, black trace, Fig. 7*G*), during light application the probability increased to almost 100% (blue trace, Fig. 7*G*) and after light-off activity returned to baseline very slowly (>100 msec). This prolonged response after light-off has been observed in several other situations for CsChrimson. For instance, at the larval neuromuscular junction (NMJ), CsChrimson drove spiking in motor neurons (monitored by muscle synaptic potentials) and exhibited a post-light response hundreds of milliseconds in duration^32^. In a second example, descending neurons DNp07 and DNp10, when expressing CsChrimson, exhibit a 100-400 msec response after light-off^33^. Finally, we tested this idea in the GF by expressing CsChrimson in the GF and illuminating the preparation at 590 nM and observed a long tail of TTM response of approximately 100 msec (Fig. 7*J*, blue trace). In brief, a variety of examples show that CsChrimson when activated produces a strong response with a long slow decay after the light is turned off (Fig 7*F-G*).

To confirm the parallel between the simulation and the *in vivo* experiment, we also developed a simulation that more closely matched the *in vivo* experiments (Fig. 7*C*). A large-scale network of the VNC, composed of leaky integrate-and-fire neuron models, was used to simulate this experiment. The GF was electrically stimulated just below threshold at 24 Hz. In these simulations, the GF only spikes in the presence of the light. Both the simulation and the optogenetic results on the living animal suggest that the Axc-to-GF connection is excitatory, consistent with the connectome and chemical predictions that the synapse is cholinergic. These results agree with our simulation’s predictions that the main effect of the optogenetic activation of the cholinergic axo-axonic neurons (Axcs) is to increase the excitability of the GF (Fig. 7*C*).

Overall, these results support our prediction that the primary effect of optogenetic activation of the cholinergic axo-axonic neurons (Axcs) is to increase the excitability of the GF. Our conclusion is that the optically activated Axcs excite the GF via their synapses in the VNC and alter the likelihood of GF action potentials.

## Discussion

By analyzing all 1,314 brain-originating descending neurons, we move beyond scattered descriptions of axo-axonic contacts and derive circuit-level design rules for presynaptic modulation. The quantitative principles that emerged within this work are: (i) extreme sparseness showing that between 0.7 to 1.2% of possible DN-to-DN, AN-to-DN, and IN-to-DN pairs form axo-axonic synapses; (ii) a tight linear relation between synaptic strength and partner multiplicity; and (iii) a small-world architecture that distributes integration rather than concentrating it in a rich-club core. Together, these features constitute a wiring grammar for axo-axonic control in the adult *Drosophila* motor system.

Axo-axonic synapses have previously been identified in the *Drosophila* brain connectome ^34–37^. Excitatory cholinergic synapses, like those investigated here, have been observed in the fly olfactory system, where projecting neurons (PN) form lateral PN-to-PN connections, similar to the DN-to-DN connections we identified^34^. Axo-axonic connections have also been reported between olfactory sensory neurons (OSNs)^36^. Bristle mechanosensory neurons (BMNs) are also shown to be axo-axonically connected^38^. These similarities suggest that axo-axonic connectivity principles might be conserved across the VNC and brains in flies, although the functional role of AACs remains unexplored.

Notably, axo-axonic wiring has revealed an extraordinarily selective, yet globally percolating schema, in contrast to, for example, the serotonin axo-axonic configuration in the hippocampus, which comprises 30%^10,39^ or in the cortex, which comprises 19%^10,40^. In the fly VNC, only ∼1 % of the possible DN-to-DN pairs form axo-axonic synapses, an order of magnitude sparser than axo-dendritic contacts in either the MANC or FAFB brain data sets^14,41^. Despite their scarcity, these connections are sufficient to incorporate every DN into a single, strongly connected network, yielding short average path lengths (∼4 hops) and thus preserving the ability for any axon to gain rapid influence over remote peers. For comparison, the entire network has an average path length of 2.8 (∼3 hops). This combination of stringent partner choice with network-wide reach resembles presynaptic inhibition motifs in mammalian spinal cord and chandelier-cell control of cortical pyramidal neurons, suggesting convergent architectural solutions across phyla^42,43^. What accounts for the sparsity of axo-axonic connections? One possibility is wiring economy: boutons on sister axons consume precious membrane and cytoplasmic resources^44^. Another is developmental lineage. Many high-degree bridge neurons derive from hemilineages that extend across neuromeres^45^, hinting that growth-cone timing and Slit-Robo–mediated midline crossing^46^ pre- pattern the opportunities for axo-axonic contact. Comparative EM work in other insects and in zebrafish larval spinal cord will clarify whether these archetypes represent a universal motor blueprint.

As previously found, rich clubs with dense interlinking of high-degree hubs are present in the fruit *Drosophila* connectome^16^. They are also widely present in vertebrates, such as in the cerebral cortex^47,48^. In contrast, the axo-axonic fly DN network revealed only a weak, transient rich-club coefficient, suggesting that presynaptic modulation in the nerve cord is decentralized. Control must therefore emerge from many mid-degree “broker” neurons (with fewer connections) and with high betweenness rather than from a minority of super-hubs. Such a layout guards against single-point failure and allows different motor modules to be gated independently, consistent with behavioral data showing that flies can blend local reflexes with whole-body maneuvers^49^.

In terms of functional implications, axo-axonic contacts positioned directly on descending axons are ideally placed to veto, amplify, or synchronize commands emanating from the brain before they fan out onto premotor interneurons and motoneurons. We have shown that Axc is amplifying GF activity. Experiments with headless animals demonstrate the existence of a second spike initiation zone (SIZ) in the VNC. We believe this is the target of the Axc synaptic modulation. Choline acetyltransferase (ChAT) immunostaining confirms that Axc type neurons are cholinergic, and optogenetic activation of these cells reliably enhances the GF’s excitability. Our simulations also demonstrate this principle: activating a cohort of eight cholinergic Axc type neurons, which exclusively target the Giant Fibers (GFs), enhances their firing.

Crucially, however, the GF is not unique; 17 other DNs receive >50 presynaptic partners that occupy bridge positions between functional modules. These neurons may be promising targets for future studies investigating the role of axo-axonic connections in precise behavior modulation. Future opto-physiological experiments probing these DNs will test whether distributed control is the norm rather than the exception.

Excitatory axo-axonic connections, exemplified by the Axc to GF projection described here, are not unique to this system and have been reported in other organisms, including the well-characterized Mauthner cells in zebrafish^12^ and dopamine cells in mice^50^. In these examples, the primary function of the excitatory axo-axonic synapse is to increase the release probability of the axon terminal they innervate. This mechanism can even bypass upstream action potentials, representing a hijacking mechanism that may have physiological roles. The findings presented in this study may contribute to a broader understanding of axo-axonic connectivity principles across species^51–55,25,56^.

In conclusion, our study presents the first comprehensive blueprint of axo-axonic connectivity within a fully reconstructed adult ventral nerve cord system. The network revealed here offers concrete parameters for next-generation models of rapid motor decision making in both invertebrates and vertebrates. By formalizing direct axon-to-axon communications, the work bridges a critical gap between connectome structure and motor function and opens new avenues for dissecting the fastest circuits in the animal kingdom.

### Limitations of study

One limitation of our study is that the connectome used in the analysis was derived from a single male specimen. New datasets are now being released that are sex-specific or can address developmental differences, such as the Female Adult Nerve Cord Connectome (FANC) and the Brain and Nerve Cord Connectome (BANC). All functional whole-network tests are *in silico*, and although experimental verification is provided through optogenetics, the large-scale network based on leaky integrate-and-fire neuron model respects synapse counts but ignores conductance dynamics and neuromodulators influences. Furthermore, EM of the MANC connectome (v1.2.1) did not detect gap-junctional coupling, suggesting that the present map under-reports electrical axo-axonic contacts that could further reshape information flow^51–55^.

Lastly, dendro-axonic synapses account for only 2% of all synapses^25,56^. Thus, we cannot rule out the possibility that some of the synapses investigated in this study are dendro-axonic, although the vast majority are axo-axonic. Nevertheless, we curated a large number of EM images (Fig. S1) to demonstrate the presence of mitochondria in axo-axonic synapses of descending neurons, suggesting that this structure can be used as a marker of axo-axonic synapses. Based on this, we found that synapses of AN08B098 onto Giant Fiber contain presynaptic mitochondria, suggesting that these synapses are axo-axonic.

We found that the number of neurons in the Gal4 line was larger than the number of neurons found in the connectome that belong to the type AN08B098. We cannot establish the source of this difference. Although we characterized two other Gal4 lines (Bloomington stock #87537 and #87664) and we found similar counts.

The long tail of response following activation of Axc in Fig 7 could be explained by multiple underlying mechanisms; local recurrent excitatory loop between GF and Axc, unknown descending neurons that feedforward to this circuit, but we are unable to identify the source of these inputs.

## STAR Methods

### KEY RESOURCES TABLE

| REAGENT or RESOURCE | SOURCE | IDENTIFIER |
| --- | --- | --- |
| **Antibodies** |  |  |
| Mouse Anti-Drosophila choline acetyltransferase | DSHB | RRID:AB_528122 |
| Mouse anti-Bruchpilot (nc82) | DSHB | RRID:AB_2314866 |
| Rabbit anti-GFP | Invitrogen | RRID:AB_221569 |
| Goat anti-rabbit secondary | Jackson ImmunoResearch | RRID:AB_2338021 |
| Goat anti-mouse Alexa Fluor 647 secondary | Jackson ImmunoResearch | RRID:AB_2338902 |
| Anti-mVenus antibody (conjugated) | MilliporeSigma | MABE1906 |
| **Bacterial and virus strains** |  |  |
| N/A | N/A | N/A |
| **Biological samples** |  |  |
| N/A | N/A | N/A |
| **Chemicals, peptides, and recombinant proteins** |  |  |
| All trans retinal 500mg | MilliporeSigma | R2500-500MG |
| Tetramethylrhodamine-labeled dextran (10,000 MW) | Invitrogen | D1817 |
| O'Dowd's Drosophila saline | <sup>57</sup> | N/A |
| Formaldehyde, Para | Fisher Scientific | T353-500 |
| Triton X-100 | MilliporeSigma | T8787-100ML |
| **Critical commercial assays** |  |  |
| N/A | N/A | N/A |
| **Deposited data** |  |  |
| N/A |  |  |
| **Experimental models: Cell lines** |  |  |
| N/A | N/A | N/A |
| **Experimental models: Organisms/strains** |  |  |
| SS25451: w[1118]; VT063643-p65.AD; VT059224-GAL4.DBD | Bloomington Drosophila Stock Center | RRID: BDSC_89495 |
| w[1118]; P{y[+t7.7] w[+mC]=20XUAS-IVS-CsChrimson.mVenus}attP2 | Bloomington Drosophila Stock Center | RRID: BDSC_55136 |
| w[*]; P{y[+t7.7] w[+mC]=20XUAS-IVS-mCD8::GFP}attP2 | Bloomington Drosophila Stock Center | RRID: BDSC_32194 |
| Control- | Bloomington Drosophila Stock Center | RRID:BDSC_ |
| **Oligonucleotides** |  |  |
| N/A | N/A | N/A |
| **Recombinant DNA** |  |  |
| N/A | N/A | N/A |
| **Software and algorithms** |  |  |
| Python | http://www.python.org/ | RRID:SCR_008394 |
| Anaconda | https://docs.anaconda.com/ | RRID:SCR_025572 |
| Jupyter Notebook | https://jupyter.org/ | RRID:SCR_018315 |
| neuprint-python API | 27 | N/A |
| NetworkX | https://networkx.org/ | RRID:SCR_016864 |
| MATLAB | https://www.mathworks.com/products/matlab.html | RRID:SCR_001622 |
| MANC | 14 | N/A |
| Inkscape | https://inkscape.org/en/ | RRID:SCR_014479 |
| Brian2 | http://briansimulator.org/ | RRID:SCR_002998 |
| NeuronBridge v3.2.6 | 58 | N/A |
| Other |  |  |
| Axon Digidata 1440A Data Acquisition System | Molecular Devices | Model 1440A |
| Clampex software (version 10 and 11) | http://www.moleculardevices.com/products/software/pclamp.html | RRID:SCR_011323 |
| Nikon AXR confocal microscope with NSPARC detector | Nikon | N/A |
| Nikon Eclipse E600FN fluorescence microscope | Nikon | N/A |
| 590 nm optogenetic LED setup | Thorlabs | M590F3; M118L02; LEDD1B |

#### Lead contact

Further information and requests for resources and reagents should be directed to and will be fulfilled by the Lead Contact, Rodrigo Pena.

#### Materials Availability

This study did not generate new unique reagents.

### EXPERIMENTAL MODELS AND SUBJECT DETAILS

#### Fly Genetics

Flies were raised at 25°C with a natural (12-hr) light cycle, and data were collected from both male and female adults 2-4 days old. The Gal4 drivers used in our study were selected through NeuronBridge^58^. Neurons of interest were identified through NeuPrint+, then identified using NeuronBridge software v3.2.6, and finally chosen based on similarity score and visual accuracy. A split-Gal4 driver specific to AN08B098 (Axc) was identified using NeuronBridge software v3.2.6 and chosen based on similarity score. SS25451 contains w[1118]; VT063643-p65.AD; VT059224-GAL4.DBD (BDSC #89495) (see Fig S4). Constructs used were 20XUAS-IVS-CsChrimson.mVenus (#55136), and 20XUAS-IVS-mCD8::GFP (BDSC #32194).

### METHOD DETAILS

#### Electrophysiology

Flies were mounted dorsal side up in dental wax after CO_2_ anesthesia. Giant Fibers were activated by extracellular stimuli applied using Grass stimulators with tungsten wire electrodes placed in the brain through the eyes, while a tungsten wire electrode was placed in the fly’s abdomen to ground the circuit ^30^. Muscle recordings were obtained using glass microelectrodes placed directly into both jump (TTM) and flight (DLM) muscles ^18,30,55,58,73^. Recording microelectrodes were filled with O’Dowd’s Drosophila saline ^74^. We do not directly monitor GF spikes but instead use TTM responses as a reporter. Extracellular stimulation of the GF pathway activates both the TTM and the DLM with a short and constant latency between them (∼0.6 ms) as described in the early recordings of this type in the 1980s^37,59^. Action potentials in the GF are indicated by synaptic potentials in both the TTM and the DLM with the constant latency between them (see Figure 7) as described in a more recent protocol for the preparation ^30^. It has long been assumed that the GF has the lowest threshold for descending neurons using this method of brain stimulation due to its large size and the presence of a spike initiation region in the brain. Recently, a spike initiation zone (SIZ) has been confirmed to be located in the brain at the dendrite-axon bifurcation ^35^. The Axon Digidata 1440A Data Acquisition System was used to digitize recordings which are then visualized using Clampex software (version 10 & 11, Molecular Devices). Muscle responses were obtained by stimulating at 24 Hz.

#### Solution preparations

Phosphate buffered saline was made from tablets purchased from Fisher BioReagents (BP2944- 100). One tablet was dissolved in double distilled water to yield a solution of 10 mM Phosphate Buffer, 2.7 mM KCl, and 137 mM NaCl. Solutions were stored at the recommended 4°C.

All trans retinal (500 mg), which was purchased from Sigma-Aldrich, is dissolved in 17.6 ml absolute ethanol to make 100 mM stock solutions. 100 µl of 100 mM stock solution is transferred to small tubes and wrapped in aluminum foil and kept in -20°C freezer (ATR should be kept away from light since it is sensitive to light and it would be degraded, thus becoming ineffective if it is exposed to light for a long time). To prepare fly food supplemented with 400 µM ATR, 5 mL fly food is dissolved in microwave. The food is left to cool down, then 20 µl of 400 µM ATR is mixed well with fly food or 20 µl of absolute ethanol is mixed with food as a control. The food vial is wrapped in aluminum foil then the food is left until it was solidified well; otherwise, the flies get stuck in the wet food. The flies are transferred from its vial to ATR containing vial and kept in a dark place (to keep the ATR from degradation) at room temperature 22-23°C.

O’Dowd saline was made by dissolving: NaCl (0.59 g); NaHCO_3_ (0.174 g); KCl (0.0224 g); MgCl_2_ (0.0814 g); CaCl_2_ (0.0148 g); NaH_2_PO_4_ (0.0138 g); Trehalose (0.1892 g); and Sucrose (1.54 g) in 100 mL double distilled water. Saline was kept in 4°C for storage or in room temperature as needed.

#### Optogenetics

We drove channelrhodopsin expression in neurons of interest using the UAS-Gal4 system. Flies expressing csChrimson (BDSC #55136) were crossed with our driver line to produce: w1118; VT063643-p65/+; 20XUAS-IVS-csChrimson.mVenus/VT059224-GAL4.DBD. Adults were placed in the dark at 50% relative humidity on standard cornmeal-molasses food, supplemented with all-trans-retinal (0.4 mM post-eclosion) for two days before testing. Light stimulation was performed for 100 ms while flies are mounted in dental wax as described above. We used a 590 nm high-power fiber-coupled LED (1.5 mW maximum light intensity, Thorlabs M590F3) with an optic fiber of diameter 200 µm and N.A. 0.22 to activate the mVenus csChrimson (M118L02). We tested various placements for the optic fiber and found the strongest activation of csChrimson to occur when the fiber was less than 1 cm from the sample and aimed at the neck. Light intensity was controlled via T-Cube LED Driver (Thorlabs LEDD1B) and kept at maximum power during testing. Each sample received electrical-only, optogenetic-only or paired opto-electrical stimulation. If Axc is stimulated only, no response is recorded, meaning that csChrimson alone is unable to elicit spikes in the GF.

We also conducted two control experiments for the optogenetic protocol (Fig. S2). The first control used flies of the same genotype as the experimental group but were not raised on food supplemented with all-trans-retinal. The second control involved flies that were raised on retinal-supplemented food but lacked the UAS-csChrimson.mVenus transgene (w1118; VT063643/+; VT059224-GAL4/VT059224-GAL4). The results are shown in Figure S2 B-C.

#### Dye-injection

The VNC and connective of young adult male flies were dissected and kept in cold O’Dowd’s saline within a Sylgard 184-filled dissection dish. Samples were immediately transferred to a Nikon Eclipse E600FN fluorescent microscope. A tetramethylrhodamine-labeled dextran dye (10,000MW, Invitrogen, D1817) solution was backfilled into a pulled glass electrode tip (∼30 MΩ resistance tip) and filled with O’Dowd’s saline. Electrodes were placed into a 3D hydraulic micromanipulator. The electrode was lowered into place, penetrating the GF axons near the neck connective. A Gettings Model 5A amplifier was used to inject positive current pulses (between 1 nA and 5 nA), with a reference electrode placed into the saline-filled dish. GFs were filled with dye for approximately 5 minutes each.

#### Immunostaining

A mouse primary antibody against ChAT (1:20, DSHB, 4B1) was used to confirm cholinergic cells and coupled to secondary Goat anti-mouse AlexaFluor 647 (1:500, 115-605-003). A mouse primary antibody against bruchpilot (1:50, DSHB, NC82) was used to label presynaptic chemical active zones and coupled to secondary Goat anti-mouse AlexaFluor 647 (1:500, 115-605-003). Primary rabbit anti-GFP (1:500, Invitrogen A11122) and secondary goat anti-rabbit (1:500, Jackson Immuno, 111-225-144) were used to amplify the fluorescence of mCD8::GFP. mVenus-tagged csChrimson. Fluorescence was amplified using a conjugated antibody (1:500, Millipore, MABE1906). Primary antibody solutions contained 3% BSA and 0.3% Triton X-100 in PBS. The secondary antibody solutions contained PBS.

Samples were dissected and processed according to the dye-injection methods above. Samples were then immediately fixed in PBS + 4% PFA for 25 minutes at room temperature (22°C). Samples were washed with PBS and placed on a rotator (60RPM) for 30 minutes. Samples were permeabilized with PBS + 0.5% Triton X-100 for 45 minutes at 22°C. Primary antibodies were added, and samples were stored at 4°C for 48 hours. Samples were then washed on a rotator with PBS for 4 hours at 22°C, changing the solution every 30 minutes. Secondary antibodies were added, and samples were stored at 4°C for 24 hours. Samples were then washed on a rotator with PBS for 4 hours at 22°C, changing the solution every 30 minutes. Samples were dehydrated with a series of ethanol steps, 50%, 70%, 90%, and 100% for 8 minutes each step. Samples were trimmed and embedded in 100% methylsalycylate, covered with cover slips, sealed, and imaged within 24 hours.

#### Confocal Microscopy

Samples were imaged on a Nikon AXR confocal using an NSPARC detector and a 60x oil immersion objective. Z-dimension step size was 194 μm for all results displayed. Laser channels used were 488 nm, 561 nm, and 640 nm. Channels were imaged in series using filters to avoid cross-excitation of close wavelengths, and 2x integration was used.

#### Image Analysis

Z-stack files were imported into Imaris software v10.2.0, and the fluorescence intensity of each channel was adjusted for best fit. GF and AN08B098 (Axc) channels were surfaced using a thresholding method that matched fluorescence in each Z dimension. Masks of each channel were created for each surface to detect colocalization of Brp or ChAT. Images were acquired in volume view and exported.

#### Computational model

To study the impact of axo-axonic projections onto DNp01 (GF), we constructed an *in-silico* model of the ventral nerve cord (VNC) by utilizing the entire MANC v1.2.1 connectome. First, we scrubbed the connectivity data by omitting connections with a total strength less than 6, then we excluded all connectivity data where the neurotransmitter type, neuron type, or neuron ID was unknown or invalid. This resulted in a total of 23,437 valid neurons and 1,152,548 connections between them, with 619,684 excitatory (cholinergic) and 532,864 inhibitory (glutamatergic or GABAergic) connections. In the fly, cholinergic receptors are excitatory, whereas glutamatergic and GABAergic receptors are inhibitory^60^.

#### Neuron model

To achieve biological plausibility while avoiding computational intractability, leaky integrate-and-fire (LIF) neuron models were used:

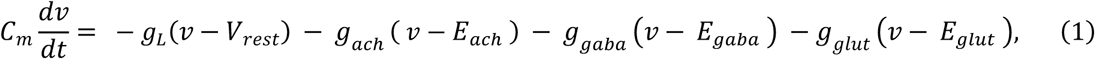

Unlike other models^61^, we employ distinct parameters for the GF compared to the rest of the neurons in the connectome, i.e., the membrane resistance. In Eq. (1), *v* is the neuron membrane potential, *g*_*L*_, *g*_*ach*_, *g*_*GABA*_, and *g*_(*lut*_ are the conductances for leak channels and acetylcholine (nicotinic), GABA (GABA_A_), and glutamate ionotropic receptors (nAChR, GABA_A_R, and GluCl, respectively). Similarly, *V*_*rest*_, *E*_*ach*_, *E*_*GABA*_, and *E*_(*lut*_ are their respective reversal potentials. Since GABA_A_R and GluR are chloride-permeable channels, their reversal potentials were obtained from chloride reversal potential^62^. *C*_*m*_ is the membrane capacitance. These are parameterized with values obtained from the literature, which investigates the biophysical properties of fruit fly synapses. Furthermore, we collected physiologically relevant values for the membrane capacitance, resistance, and leak reversal potential. Simulations of these neurons were made using the BRIAN2 simulator ^63^.

#### Synapse model

For each of the synaptic currents (*x* = *ach*, *gaba*, *glut*), the conductance obeys

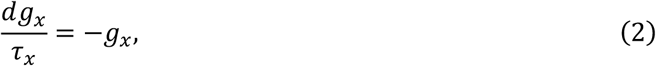

which increases by *J*_*x*_ upon a presynaptic spike. The list of parameters is included in Table 1.

**Table 1.** List of parameters used in the connectome simulation of the VNC in this paper. For each of our simulations, we ran the entire VNC network with a step-size of 0.1 ms during 2000 ms. Furthermore, for each simulation, each neuron in the VNC network was provided with an excitatory Poisson input of 30 Hz at a synaptic conductance of 5 nS to simulate typical endogenous activity driven by neurons in the fly brain. During stimulation experiments, neurons DNp01, DNg02, and/or DNd03 were stimulated with additional Poisson input at 10 Hz at a synaptic conductance of 40 nS. In Figure 8, the same network was used, but the GF was electrically stimulated just below threshold with a Poisson-distributed stimulation at 24 Hz.

| Parameter | Symbol | Value | Ref. |
| --- | --- | --- | --- |
| Acetylcholine receptor (nAChR) conductance | $J_{ach}$ | 0.27 nS | <sup>64</sup> |
| nAChR decay time | $\tau_{ach}$ | 1.1 ms | <sup>64</sup> |
| nAChR reversal potential | $E_{ach}$ | 0 mV | <sup>65</sup> |
| GABA receptor (GABA<sub>A</sub>R) conductance | $J_{GABA}$ | 0.8 nS | Estimated |
| GABA<sub>A</sub>R decay time | $\tau_{GABA}$ | 5.4 ms | <sup>66</sup> |
| GABA<sub>A</sub>R reversal potential | $E_{GABA}$ | -60 mV | <sup>60,62</sup> |
| Glutamate receptor (GluCl) conductance | $J_{glut}$ | 0.8 nS | Estimated |
| GluCl decay time | $\tau_{glut}$ | 5 ms | <sup>67</sup> |
| GluCl reversal potential | $E_{glut}$ | -60 mV | <sup>62</sup> |
| Membrane Resistance (GF) | $R_m$ | 25 MOhm | <sup>64</sup> |
| Membrane Resistance (other neurons) | $R_m$ | 300 MOhm | <sup>68</sup> |
| Resting membrane potential | $V_{rest}$ | -55 mV | 61,68 |
| Action potential threshold | $V_{th}$ | -40 mV | 68 |
| Reset potential | $V_{reset}$ | -55 mV | Estimated |
| Refractory period | $\tau_{ref}$ | 2 ms | Estimated |
| Membrane capacitance | $C_m$ | 72 pF | 68 |

### QUANTIFICATION AND STATISTICAL ANALYSIS

#### Data acquisition and analysis

Data for VNC neurons used in the study came from the publicly available *Drosophila* male adult nerve cord connectome^14^. NeuPrint+ API^26,28^ access is granted to registered users, and Python accessibility is maintained through the neuPrint-python package^27^. The latest *Drosophila* MANC data can be accessed here: https://www.janelia.org/project-team/flyem/manc-connectome.

#### Connectivity analysis

We retrieved all simple synaptic connections using neuprint.fetch_simple_connections^26–28^ separately for: outputs (presynaptic AN, IN, and DN as presynaptic) and Inputs (postsynaptic DN). Connections were summed by partner neuron and filtered to those with total strength ≥ 6. For each DN, we then counted the number of distinct AN, IN, and DN partners in both the output and input directions.

#### Network analysis

Network analysis was performed using Python with the NetworkX^69^ library to extract quantitative measures of connectivity. From the structural connectivity matrix, we calculated a comprehensive set of graph-theoretical metrics to characterize the network topology. These included basic properties (number of nodes and edges, network density), integration measures (average path length, diameter), segregation indices (clustering coefficient, modularity with community detection using the Louvain algorithm), centrality metrics (degree, betweenness, closeness, and eigenvector centrality), and organizational principles (degree assortativity, rich club coefficient).

The rich club analysis was specifically conducted across multiple degree thresholds to identify potential hierarchical organization among high-degree hub regions. All weighted graph measures incorporated the synaptic strengths from the original connectivity matrix. This multimodal graph-theoretical approach allowed for a detailed characterization of both local and global properties of the network’s structural connectome, revealing important aspects of neural communication efficiency, functional segregation, and network resilience.

#### Statistical and visual analysis

SWC analysis was conducted in Python using the Navis client^70^. We stored morphology and input/output counts using the Pandas library. Relationships between morphology metrics (cable length, branch points, node count) and partner counts were assessed with Pearson correlation and linear regression (scipy.stats.pearsonr, linregress). Results were visualized with Matplotlib^71^. Network-style connection diagrams were rendered interactively with ipycytoscape.

#### Software and libraries

NeuPrint+ analyses were performed in Python 3.12.4 using the neuprint-python API. We used the Janelia NeuPrint client (neuprint), and Navis (v1.10.0) for data access and network building; data manipulation and grouping were done with pandas v2.2.2 and NumPy v1.26.4; graph-based operations with NetworkX v3.2.1; interactive visualization with ipycytoscape v1.3.3 and ipywidgets v8.1.5; plotting and PDF export with Matplotlib v3.10.1; and statistical analyses (Pearson’s r, linear regression) with SciPy^72^ v1.13.1. Auxiliary parsing and utility functions relied on Python’s built-in json, re, itertools, and warnings modules.

## Acknowledgments

R.P. was funded by an IBRO Collaborative Research Grants. The authors thank the FAU Jupiter Life Sciences Initiative (JLSI) for funding.

## Author contributions

C.C.C. and R.P. were responsible for the conceptualization of the work. C.C.C., J.L., T.R., D.S., S.J., and C.S. analyzed the data and made the figures. R.P. worked on the graph theoretical measures. D.S. performed and analyzed the electrophysiology recordings. S.J. and C.S. performed immunofluorescence. K.R. maintained fly stocks and set up experiment crosses. T.R. built and ran the network simulations. R.P., C.S., and R.M. supervised the research and secured funding. All authors participated in the writing, reviewed, and approved the manuscript.

## HIGHLIGHTS

- Axo-axonic connections made with descending neurons (DNs) are rare in the VNC.
- There is a linear correlation between the number of synapses and the number of partners a neuron has.
- Axo-axonic neurons form exclusive connections with the Giant Fibers.
- Axo-axonic neuron activity influences DN excitability *in silico* and *in vivo*.

