## Supporting Material for "The Drosophila connectome reveals Axo-Axonic Synapses on Descending Neurons"

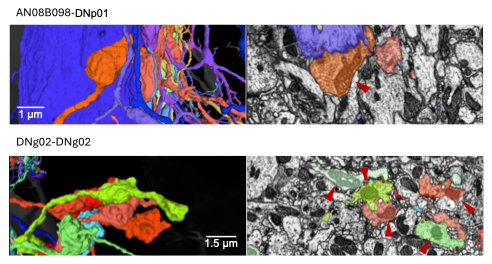

**Figure S1.** Top: Electron microscopy reconstruction (left) and single section (right) showing the synapses from AN08B098 neurons to the GF (in purple). Bottom: Electron microscopy reconstruction (left) and single section (right) showing axo-axonic synapses between DNg02 neurons. Note that presynaptic sites correspond to axonal blebs containing mitochondria closer than 3 μm.

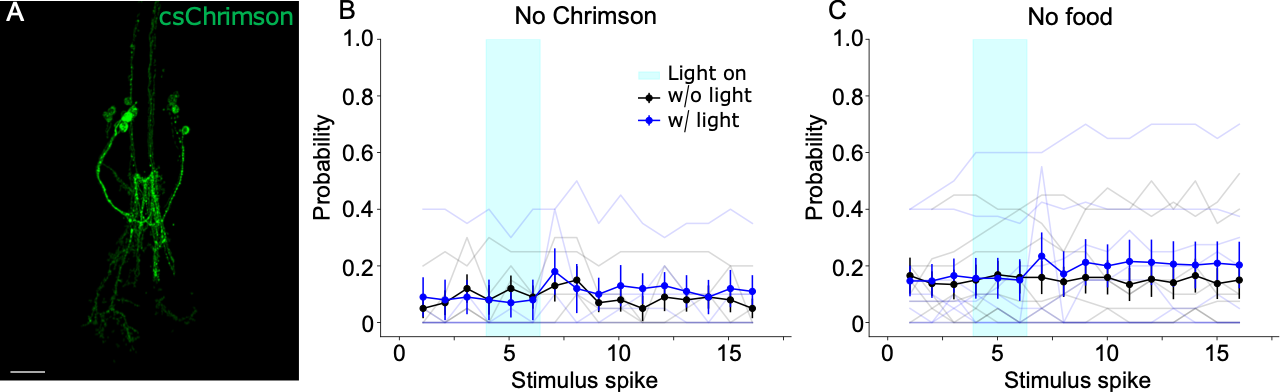

**Figure S2.** (*A*) Immunofluorescence of mVenus-tagged CsChrimson. CsChrimson was expressed along the axons and within the somas of the AN08B098 population. Scale bar represents 20 μm. (B) Electrical recordings from the jump muscle of flies lacking CsChrimson expression (n = 5 flies). (*C*) Electrical recordings from the jump muscle of flies expressing CsChrimson in AN08B098 neurons, but not provided with food supplemented with all-trans-retinal (n = 8 flies). No significant differences were observed between treatment groups.

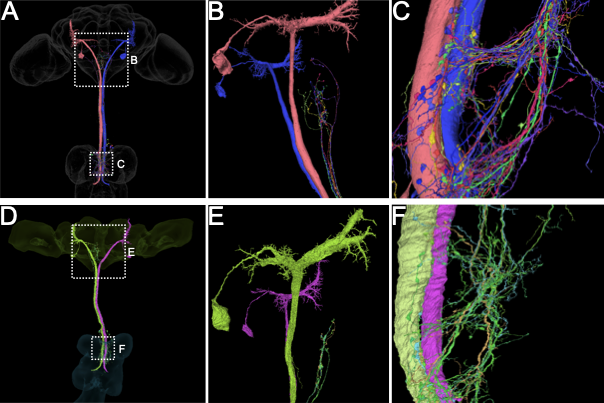

**Figure S3. Axoaxonic connections between the Giant Fibers and AN08B098 only occur in the ventral nerve cord, not in the brain.** (*A*-*C*) Reconstructions of the GFs and AN08B098 neurons from both male-cns:v0.9 (top row) and BANC (*D*-*F*, bottom row). (A) Morphological reconstruction in the brain, connective, and VNC, (B) only in the brain, (C), and only in the VNC. (D) Morphological reconstruction in the brain, connective, and VNC, (E) only in the brain, (F), and only in the VNC. Both datasets show synapses occurring between the two in the VNC, but fail to show any monosynaptic connectivity in the brain. Then, we performed pathway analysis between the two neurons using neuprint.queries.fetch_paths (see methods) to search for polysynaptic connections. This function identifies the start and endpoints of neurons and recursively traces all paths between them. In male-cns:v0.9, we manually verified all paths between AN08B098 and the GFs (DNp01) with one intermediate neuron between the two and found no connections between the two occurring in the brain. We also checked cases with 3 intermediate neurons between the two: 1) AN08B098 (162384) –12 connections--> DNg40 (10361) --14 connections --> DNp01 (10010), 2) AN08B098 (162487) –18 connections --> GNG103 (10202) –14 connections --> DNp01 (10001). Neither pathway dominates the connections to the GFs, and they are highly unlikely to drive an action potential in them.

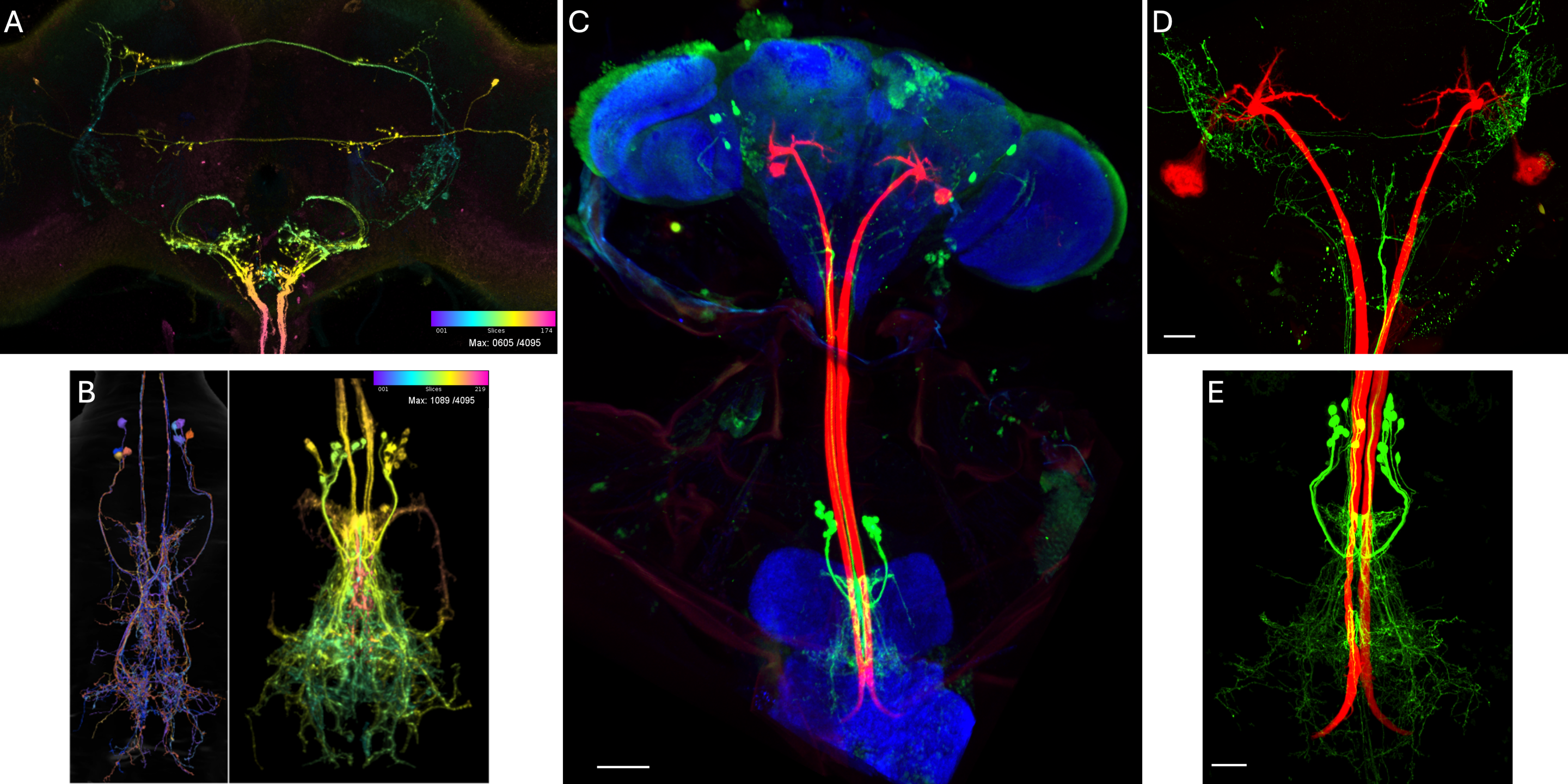

**Figure S4. Comparison of the SS25451 driver expression with EM reconstructions of AN08B098-type neurons from the MANC connectome and confocal data from the entire CNS.** (*A*) The SS25451 split-Gal4 line expresses strongly in AN08B098 neurons in the brain, as well as in additional cells, including the LCe07 visual projection neurons (images from NeuronBridge). Using Neuprint+, we did not find any monosynaptic connections between the additional neurons expressing this split-Gal4 and the GFs in the brain. (*B*) The SS25451 driver line was identified using NeuPrint+ links to NeuronBridge, and we assessed it as the best candidate for driving the expression of gene constructs in the entire population of AN08B098 neurons. While the number and location of AN08B098 neurons identified in connectomes vary between datasets, we validated their connectivity to the Giant Fibers and judged this driver to be the best for our experiments. (*C*) 20x 3D confocal stack showing SS25451 driver expression in the entire CNS. SS25451 (green) is driving UAS-mCD8::GFP, giant fibers (red) are dye-filled with tetramethylrhodamine, and neuropil (blue) is stained with NC82. Scale bar is 50 µm. (*D*) 60x 3D confocal stack showing GFs (red), and SS25451 (green). We confirmed that the split-Gal4 does not make monosynaptic connections to the GFs in the brain within our confocal slices, where we were unable to detect colocalization between the red and green fluorescence. Scale bar represents 20 µm. (*E*) 60x 3D confocal stack showing GFs (red), and SS25451 (green). As observed in SS25451 in NeuronBridge, our confocal data shows that the only cells expressing this split-Gal4 are the AN08B098 cells. Scale bar represents 20 µm.
